# The Human Eye Transcriptome Atlas: A Searchable Comparative Transcriptome Database for Healthy and Diseased Human Eye Tissue

**DOI:** 10.1101/2021.11.04.467318

**Authors:** Julian Wolf, Stefaniya Boneva, Anja Schlecht, Thabo Lapp, Claudia Auw-Haedrich, Wolf Lagrèze, Hansjürgen Agostini, Thomas Reinhard, Günther Schlunck, Clemens Lange

## Abstract

The applications of deep sequencing technologies in life science research and clinical diagnostics have increased rapidly over the last decade. Although fast algorithms for data processing exist, intuitive, portable solutions for data analysis are still rare. For this purpose, we developed a web-based transcriptome database, which provides a platform-independent, intuitive solution to easily explore and compare ocular gene expression of 100 diseased and healthy human tissue samples from 15 different tissue types collected at the Eye Center of the University of Freiburg. To ensure comparability of expression between different tissues, reads were normalized across all 100 samples. Differentially expressed genes were calculated between each tissue type to determine tissue-specific genes. Unsupervised analysis of all 100 samples revealed an accurate clustering according to different tissue types. Cluster analysis based on known cell type-specific marker genes allowed differentiation of respective tissues. Several tissue-specific marker genes were identified. These genes were involved in tissue- or disease-specific processes, such as myelination for the optic nerve, visual perception for retina, keratinocyte differentiation for conjunctival carcinoma, as well as endothelial cell migration for choroidal neovascularization membranes. The results are accessible at the Human Eye Transcriptome Atlas website at https://www.eye-transcriptome.com. In summary, this searchable transcriptome database enables easy exploration of ocular gene expression in healthy and diseased human ocular tissues without bioinformatics expertise. Thus, it provides rapid access to detailed insights into the molecular mechanisms of various ocular tissues and diseases, as well as the rapid retrieval of potential new diagnostic and therapeutic targets.

## INTRODUCTION

Gene expression analysis using high-throughput RNA sequencing provides important insights into the functional state of a cell or a tissue and has helped to improve the understanding of many diseases. Databases such as the Genotype-Tissue Expression project ^1^ or the Cancer Genome Atlas ^2^ offer a wide range of transcriptome datasets of several healthy and diseased human tissue types. However, with the exception of uveal melanoma ^2^, no ocular tissues are included in these databases. To address this challenge, Swamy et al. ^3^ developed a transcriptome database, which allows to explore gene expression in a large cohort of selected healthy human eye tissues, including retina, retinal pigment epithelium, cornea and lens. Other published ocular transcriptome databases are focused on human retinal specimens ^4^, or include murine lens ^5,6^ or retinal ^6^ samples only. Thus, none of the existing databases provides data on diseased human ocular tissue, which would be an important prerequisite to identify relevant molecular mediators, disease-related signaling pathways and novel diagnostic and prognostic markers ^7^.

For this reason, we have developed the Human Eye Transcriptome Atlas (https://www.eye-transcriptome.com), which currently includes 100 human diseased and healthy ocular samples. Unlike previously existing databases, our atlas provides data on additional tissue types such as conjunctival melanoma, squamous cell carcinoma, papilloma, pterygium, eyelid, lacrimal gland, as well as choroidal neovascular membranes, choroid/RPE, optic nerve, retinal microglia and hyalocytes. The database allows the user to easily explore and compare ocular gene expression in diseased and healthy eye tissue without bioinformatics expertise. This provides easy and rapid access to obtain detailed insights into the molecular mechanisms and functional state of various ocular tissues and diseases and allows for the rapid retrieval of potential new diagnostic and therapeutic targets for various eye diseases in the future.

## RESULTS

### Included diseased and healthy eye tissues

The Human Eye Transcriptome Atlas includes the transcriptional profile of a total of 100 human diseased and healthy eye specimens comprising 12 conjunctival melanoma ^7^, 7 conjunctival squamous cell carcinoma (SCC) ^8^, 7 papilloma ^8^, 8 pterygia and 4 choroidal neovascularization membranes ^12^, as well as 11 healthy conjunctiva ^7,8^, 3 cornea, 6 eyelid, 8 lacrimal gland, 4 optic nerve, 6 retina (3 centre and 3 periphery), 4 choroid/retinal pigment epithelium ^12^, 6 retinal microglia and 14 hyalocyte ^9^samples. Details about the included specimens and demographic data of all patients are summarized in Table 1.

**Table 1:** Healthy and diseased human eye tissues included in the database. Data is shown as mean (standard deviation) or as absolute numbers. CNV: choroidal neovascularization, FFPE: Formalin fixation and paraffin embedding, RPE: retinal pigment epithelium, SCC: squamous cell carcinoma.

|  | Tissue | n | Age at surgery (y) | Sex (m/f) | FFPE/unfixed |
| --- | --- | --- | --- | --- | --- |
| Diseased | Conjunctival melanoma <sup>7</sup> | 12 | 58.9 (17.9) | 3/9 | 12/0 |
|  | Conjunctival SCC <sup>8</sup> | 7 | 69.6 (13.3) | 5/2 | 7/0 |
|  | Conjunctival papilloma <sup>8</sup> | 7 | 37.1 (24.3) | 3/4 | 7/0 |
|  | Pterygium | 8 | 57.6 (8.5) | 6/2 | 8/0 |
|  | CNV membrane <sup>12</sup> | 4 | 79.6 (7.6) | 1/3 | 4/0 |
| Healthy | Conjunctiva <sup>7,8</sup> | 11 | 56.6 (7.5) | 9/2 | 8/3 |
|  | Cornea | 3 | 69.6 (13.2) | 2/1 | 3/0 |
|  | Eye lid | 6 | 79.3 (13.4) | 4/2 | 6/0 |
|  | Lacrimal gland | 8 | 47.7 (18.5) | 3/5 | 8/0 |
|  | Optic nerve | 4 | 54.8 (4.1) | 3/1 | 4/0 |
|  | Retina periphery | 3 | 68.5 (21.1) | 1/2 | 3/0 |
|  | Retina centre | 3 | 54.8 (5.0) | 2/1 | 3/0 |
|  | Choroid/RPE <sup>12</sup> | 4 | 70.4 (3.8) | 3/1 | 4/0 |
|  | Retinal microglia | 6 | 65.4 (17.1) | 4/2 | 0/6 |
|  | Hyalocytes <sup>9</sup> | 14 | 70.8 (12.5) | 8/6 | 0/14 |
|  | Total | 100 | 61.8 (17.3) | 57/43 | 77/23 |

### Unsupervised transcriptomic analysis

The unsupervised analysis of all 100 samples revealed an accurate clustering according to different tissue types (Fig. 1). As expected, the highest amount of variation was observed between the cell type-specific samples (retinal microglia as well as hyalocytes) and the remaining whole tissue samples, which are composed of a mixture of various cell types.

**Figure 1:**
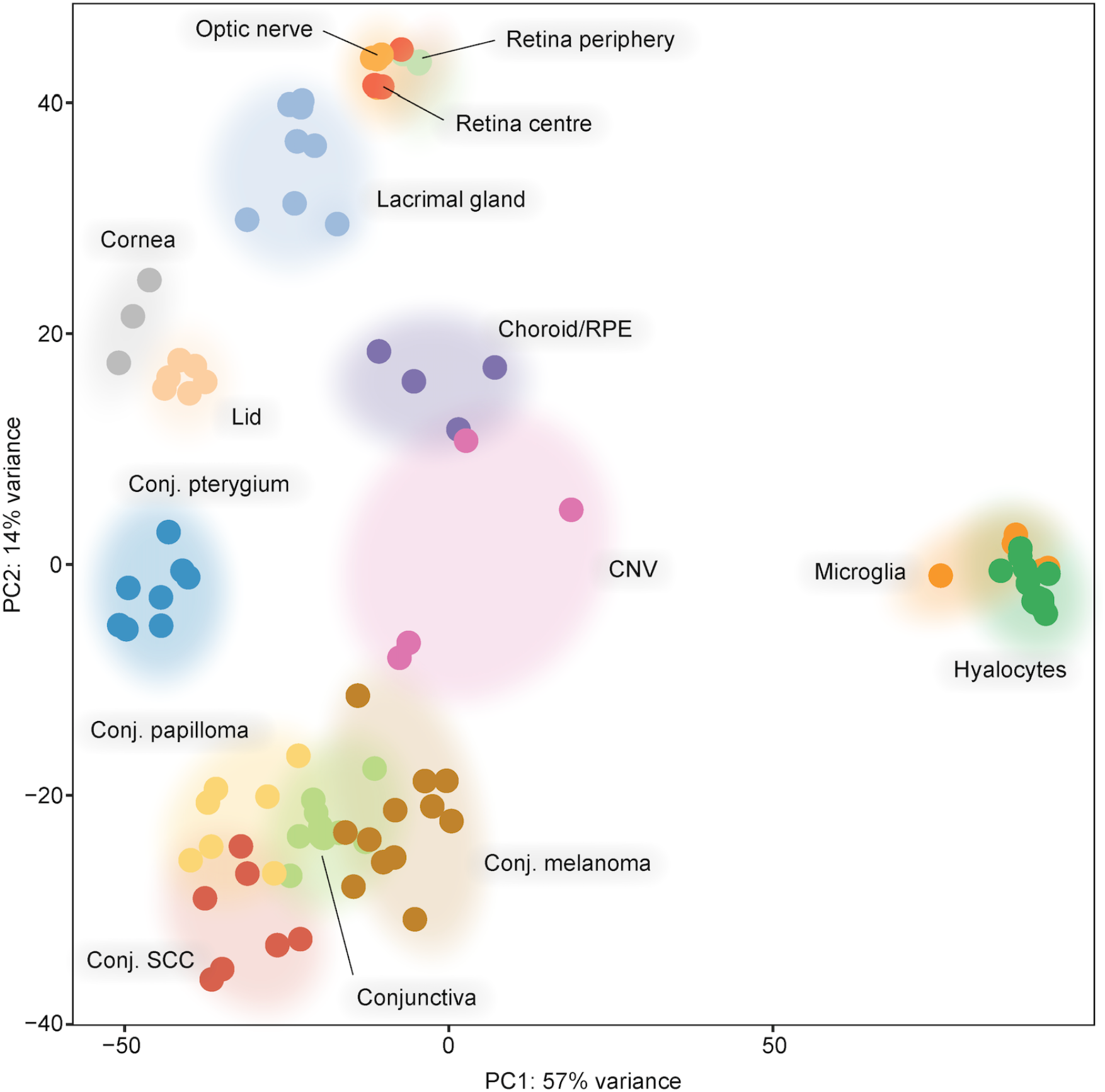
Principle component analysis (PCA) demonstrates clustering of samples by tissue type. Each dot represents one sample. CNV: choroidal neovascularization, Conj.: conjunctival, PC: Principle component, RPE: retinal pigment epithelium, SCC: squamous cell carcinoma.

### Tissue-specific genes

Tissue-specific genes were identified by calculating differentially expressed genes between the tissue type of interest and the remaining samples in a first step. Only transcripts with an adjusted *p*-value < 0.05 were considered for further analysis. In a second step, genes for which the 25^th^ percentile of expression in the tissue type of interest was higher than the 90^th^ percentile of expression in all other tissues were considered tissue-specific genes (see methods for details). The expression profile of the most specific genes for each tissue type is visualized in the heatmap in Figure 2A. Among the most specific marker genes, there were *KRT6B* (keratin 6B) and *KRT75* for conjunctival SCC, *MIA* (Melanoma Inhibitory Activity) and *S100A1* (S100 calcium binding protein A1) for conjunctival melanoma, *ANGPTL7* (Angiopoietin Like 7) and *COL8A2* (Collagen Type VIII Alpha 2 Chain) for cornea, *LCN1* (lipocalin 1) and *LTF* (Lactotransferrin) for lacrimal gland, *PLP1* (Proteolipid Protein 1) and *AQP4* (Aquaporin 4) for optic nerve, *RHO* (rhodopsin) and *PDE6G* (phosphodiesterase 6G) for retina, as well as *ACTN4* (Actinin Alpha 4) and *PLXND1* (Plexin D1) for CNV membranes (Fig. 2A). Gene ontology (GO) analysis revealed specific genes associated with disease-relevant biological processes (Fig. 2B and Supplement Fig. 1), such as chromatin organization and angiogenesis for conjunctival melanoma, keratinocyte differentiation and epidermis development for conjunctival SCC, glandular epithelial cell development for healthy conjunctiva, extracellular matrix organization and regulation of cell-matrix adhesion for cornea, lipid biosynthetic process for eyelid, glia cell differentiation and myelination for optic nerve, endothelial cell migration and actin filament organization for CNV membranes as well as visual perception and phototransduction for retina (Fig. 2B and Supplement Fig. 1).

**Figure 2:**
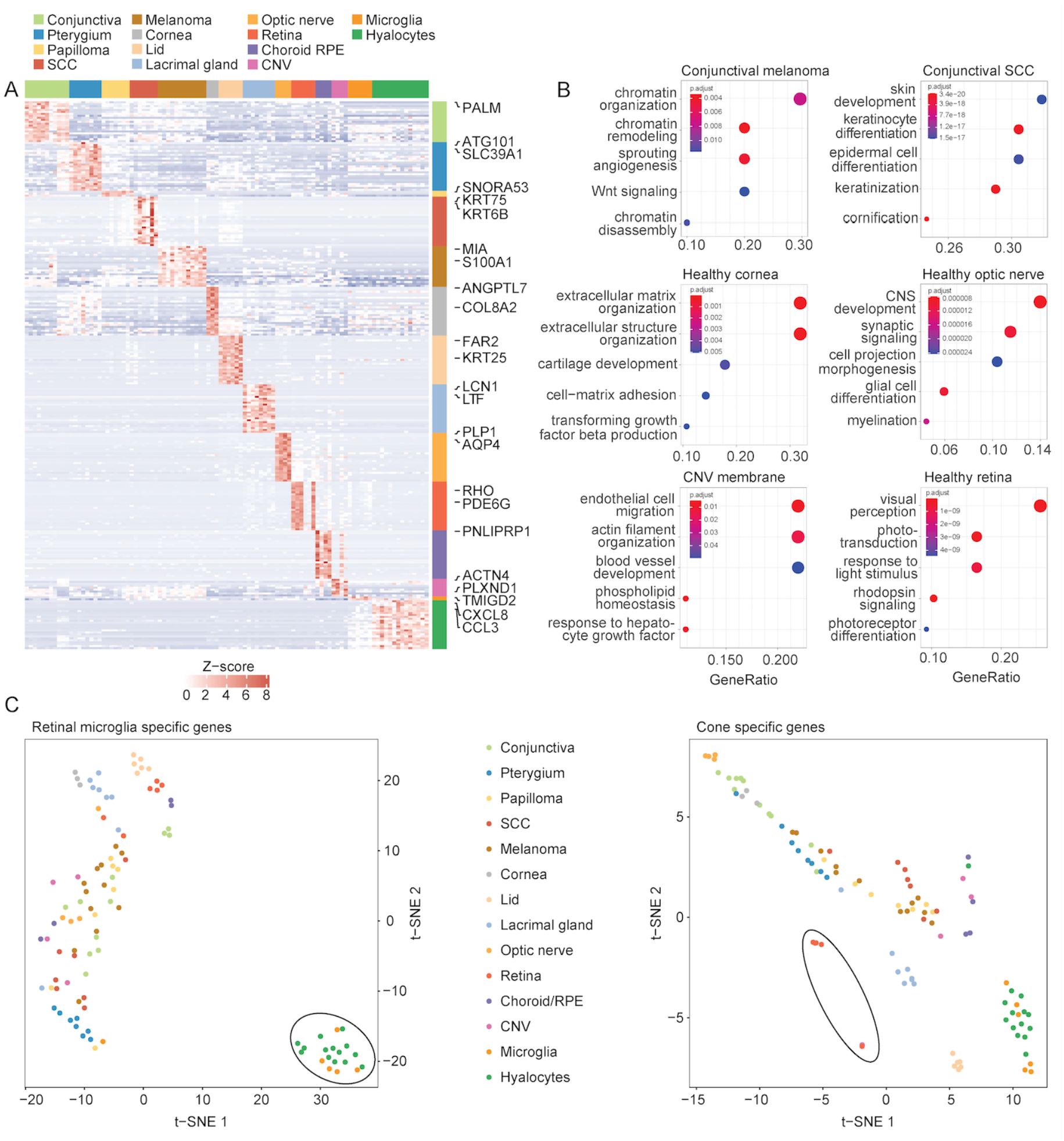
Tissue-specific genes. (A): Heatmap visualizing tissue-specific genes in the Human Eye Transcriptome Atlas (maximum of 25 genes for each tissue). Each column represents one sample and each row represents one gene (see colored legend for different tissue types). Selected specific marker genes are labeled on the right. The z-score represents a gene’s expression in relation to its mean expression in standard deviation units (red: upregulation, blue: downregulation). (B): Gene ontology (GO) analysis of all specific genes in conjunctival melanoma, conjunctival squamous cell carcinoma, cornea, optic nerve, choroidal neovascularization and retina (see Supplement Figure 1 for remaining tissues). The top five biological processes, which the specific genes are involved in, are illustrated in the dot plots. The size of the dots represents the number of associated genes (count). The adjusted p-value of each GO term is shown by color. The gene ratio describes the ratio of the count to the number of all specific genes. (C-D): Validation of the Human Eye Transcriptome Atlas based on known cell type-specific marker genes for retinal microglia (C) and cones (D) using t-distributed stochastic neighbor embedding (t-SNE) plots. Hyalocytes and retinal microglia clearly separated from all other samples based on the expression of 100 retinal microglia specific marker genes derived from single cell RNA sequencing (C, black circle). Additionally, the retinal specimens in the Human Eye Transcriptome Atlas can be distinguished from other samples based on cone specific marker genes (D, black circle). Each dot represents one sample. Color represents different tissue types (see legend between C and D).

Additionally, transcriptional profiles of samples in the Human Eye Transcriptome Atlas were validated using known cell type-specific marker genes for retinal microglia and cones derived from single cell RNA sequencing studies ^22^ (Fig. 2 C-D). Cluster analysis based on the expression of 100 most specific retinal microglia marker genes revealed a distinct clustering of retinal microglia and hyalocytes compared to all other samples (Fig. 2C). Similarly, retinal samples were differentiated from other specimens based on the expression of cone-specific marker genes (Fig. 2D).

### Human Eye Transcriptome Atlas website

We designed the Human Eye Transcriptome Atlas website (https://www.eye-transcriptome.com) to allow a user to easily explore gene expression data of various diseased and healthy human eye tissues, quality-assured through processing at a single center, without bioinformatics expertise nor the need for time-consuming bioinformatics pipelines.

The core functionality is a search engine, which provides the user with a tool to search for the expression of any Hugo Gene Nomenclature Committee (HGNC) ^24^ gene symbol, which is visualized using boxplots (Fig. 3A). The user can select tissue types by categories (e.g. all tissues from the anterior or posterior segment of the eye as well as healthy or diseased tissue) or individually. To enable the use of expression data for the user’s own research projects, normalized reads of a gene in the selected datasets can be downloaded by clicking on the “Download CSV”-button below the plot. For more information about the selected gene, the user can click on the title of the plot, which links to the corresponding gene’s page in the GeneCards database ^25^ (Fig. 3A). Another key function is the exploration of tissue-specific marker genes throughout each included dataset (Fig. 3B), which can be visualized using modified volcano-plots (Fig. 3B). Each dot represents one tissue-specific gene. By moving the mouse to a specific point, the gene’s symbol is displayed. A click on the gene visualizes its expression in all samples as illustrated in Fig. 3C.

**Figure 3:**
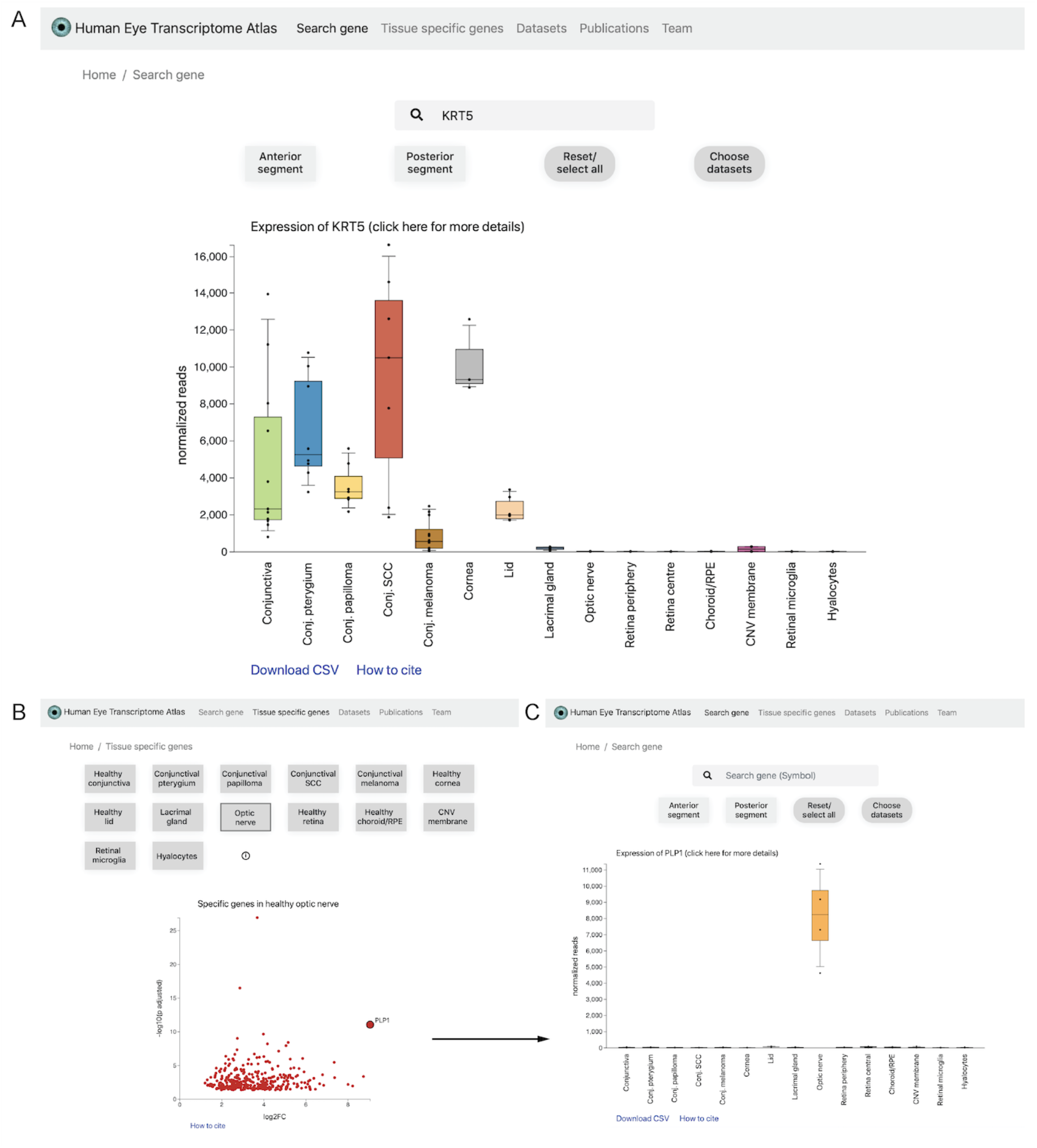
Human Eye Transcriptome Atlas website (https://www.eye-transcriptome.com). (A): Search gene function: The user can easily search the Human Eye Transcriptome Atlas for any HGNC gene symbol without bioinformatics expertise. Tissues can be selected according to the user’s preferences or by selecting categories (e.g. all tissues from the anterior or posterior segment of the eye as well as healthy or diseased tissue). The expression of the queried gene in all selected datasets is visualized using boxplots. The user can download the expression data shown in the boxplots by clicking on the “Download CSV”-button below the plot. For more information about the selected gene, the user can click on the title of the plot, which links to the corresponding gene’s page in the GeneCards database. (B): Tissue-specific genes of all included datasets can be visualized using modified volcano-plots. The selected example shows the expression of proteolipid protein 1 (PLP1), which is a predominant component of myelin in central nerves. Each dot represents one tissue-specific gene. The x-axis shows the log2 fold change between the selected tissue and all other tissues. The y-axis represents the adjusted p-value (log10-scaled). Moving the mouse to each point displays the gene’s symbol (e.g. *PLP1*). A click on one gene visualizes its expression in all samples (C, *PLP1*). HGNC: Hugo Gene Nomenclature Committee ^24^.

## DISCUSSION

Transcriptional profiling of diseased human eye tissue allows to obtain additional information about disease-relevant signaling pathways as well as about new diagnostic and prognostic markers ^7,8,12^. Since the analysis of raw sequencing data requires bioinformatics expertise, we have developed the Human Eye Transcriptome Atlas (https://www.eye-transcriptome.com) which allows the user to easily explore ocular gene expression in 100 diseased and healthy eye tissue samples without bioinformatics knowledge. Since existing ocular transcriptome databases only contain healthy specimens of selected tissue types ^3-6^, the Human Eye Transcriptome Atlas additionally provides several diseased ocular entities, such as conjunctival melanoma, squamous cell carcinoma, papilloma, pterygium and choroidal neovascular membranes as well as some healthy entities not yet available in any searchable database, such as lacrimal gland, eyelid, optic nerve, retinal microglia and hyalocytes.

To ensure high quality, tissue specificity and comparability between samples, only specimens collected at our institution were included without considering publicly available transcriptome data from third parties. All histological diagnoses were confirmed by two experienced ophthalmic pathologists and only specimens with a clear histological diagnosis were sequenced. This ensures a high quality and tissue specificity of transcriptional profiles, as demonstrated by accurate clustering according to tissue type, association of tissue-specific marker genes in disease-relevant biological processes and the validation based on known cell type-specific marker genes. Considering that post-mortem tissue samples may be subject to rapid RNA degradation as a result of an extended death to preservation time ^12,26,27^, no human donor eyes were included in this study. Instead, only samples which were immediately processed after surgical excision were included. In addition, the same sequencing protocol was used for all FFPE samples ^8^, which minimizes technical variability between different datasets. Retinal microglia and hyalocytes, in contrast, were sequenced using a different protocol, which is optimized for unfixed samples ^9^. This may have added some technical variability. However, a recent study from our group compared both protocols and revealed no substantial differences in expressed gene patterns ^8^, which indicates a low probability of a significant batch effect. To ensure comparability of expression between different datasets, all 100 samples were included in one DESeq2 ^17^ model to calculate normalized reads. It should be noted that this may lead to discrepancies in relative expression values compared to the original publications of individual datasets and that, in addition, expression values may change in the future when new datasets will be added.

The present study also identified several tissue-specific marker genes for 14 different healthy and diseased tissue types of the anterior and posterior segment of the human eye. These genes were involved in disease- or tissue-specific processes, such as keratinocyte differentiation for conjunctival SCC, chromatin organization for conjunctival melanoma, glandular epithelial cell development for healthy conjunctiva, extracellular matrix organization for cornea, lipid biosynthetic process for eyelid, myelination for optic nerve, endothelial cell migration for CNV membranes as well as visual perception for retina, which underlines the biological relevance of these markers. Some of the identified genes have already been described as markers for a related non-ocular entity, such as *KRT6B* for SCC of the lung ^28^, as well as *MIA* and *S100A1* for skin melanoma ^29,30^. *PLXND1*, which was one of the CNV specific genes, was found to be upregulated in endothelial cells during tumor angiogenesis ^31^. In addition, among the identified tissue-specific genes, there were several factors, which have already been reported in association with tissue-specific functions, among them *ANGPTL7*, playing a major role in maintaining corneal avascularity and transparency ^32^, *COL8A2*, being the main collagen type in Descemet’s membrane ^33^, with mutations being associated to Fuchs’ corneal dystrophy ^34^, as well as *PLP1*, being the predominant component of myelin ^35^ and *AQP4*, being associated to neuromyelitis optica ^36^.

In summary, the present study adds a searchable and comparative transcriptome database for healthy and diseased ocular tissue, which allows the user to easily explore ocular gene expression without bioinformatics expertise. The website thus lays the foundation for a user-friendly transcriptome library, which will be regularly updated and expanded with new datasets from our institution. This will enable scientists to quickly test hypothesis and identify potential new diagnostic and therapeutic targets for various ocular diseases in the future.

## METHODS

### Tissue specimens

A total of 100 different healthy and diseased human eye tissue samples, which were obtained at the Eye Center of the University of Freiburg between 1996 and 2020, were retrospectively included in this study. All methods were carried out in accordance with relevant guidelines and regulations. Ethics approval was granted from the Ethics Committee of the Albert-Ludwigs-University Freiburg, Germany (approval numbers 17/17 and 481/19).

### Tissue processing

Resected tissue was processed as previously described ^8-10^. In brief, samples were either processed unfixed ^9^ or formalin-fixed and embedded in paraffin ^8^ immediately after surgery according to routine protocols. Histological diagnoses were confirmed by two experienced ophthalmic pathologists. For cell analysis, retinal microglia or vitreous hyalocytes were FACS sorted and isolated according to a standardized protocol ^9^. A detailed overview of the analyzed samples, the associated publications and the respective tissue processing manner can be found in Table 1.

### RNA Sequencing

For FFPE samples, RNA sequencing was performed using massive analysis of cDNA ends (MACE), a 3’-RNA sequencing method, as previously described ^7,8,11,12^, which allows sequencing of FFPE samples with high accuracy ^8^. Briefly, barcoded libraries comprising unique molecule identifiers were sequenced on the NextSeq 500 (Illumina) with 1 × 75 bp. PCR bias was removed using unique molecular identifiers. For unfixed samples, RNA sequencing was performed as previously described ^8,9^.

### Bioinformatics

Sequencing data (fastq files) were uploaded to and analyzed on the Galaxy web platform (usegalaxy.eu) ^13^. Quality control was performed with *FastQC Galaxy Version 0*.*72* (http://www.bioinformatics.babraham.ac.uk/projects/fastqc/ last access on 08/10/2020). Reads were mapped to the human reference genome (Gencode, release 34) with *RNA STAR Galaxy Version 2*.*7*.*5b* ^*14*^ with default parameters using the Gencode annotation file (Gencode, release 34, https://www.gencodegenes.org/human/releases.html). Reads mapped to the human reference genome were counted using *featureCounts Galaxy Version 1*.*6*.*4* ^15^ with default parameters using the aforementioned annotation file. The output of featureCounts was imported to RStudio (Version 1.2.1335, R Version 3.5.3). Gene symbols and gene types were determined based on ENSEMBL release 100 (Human genes, download on 05/25/2020) ^16^. Genes with zero reads in all samples were removed from analysis. Principal component analysis (PCA) ^17^ was applied to check for potential batch effects, which were removed by the *limma* function *removeBatchEffect* ^*18*^ and by consideration within the linear model of DESeq2 Version 1.22.2 ^17^. To ensure comparability of expression between different tissues, all 100 samples were included in in one model of DESeq2 ^17^ (default parameters). Note that normalized reads might change in the future when new datasets will be added. Heatmaps were created using the R package *ComplexHeatmap 1*.*20*.*0* ^*19*^. Other data visualization was performed using the *ggplot2* package ^20^. Gene enrichment analysis and its visualization were done using the R package *clusterProfiler 3*.*10*.*1* ^*21*^.

Genes differentially expressed between the samples of the respective tissue and the remaining samples were calculated using DESeq2 ^17^ to determine tissue-specific genes (Benjamini-Hochberg adjusted p-values). Transcripts were filtered for adjusted *p*-value < 0.05. Additionally, the 25^th^ and the 90^th^ percentile of expression of each gene in each tissue type were calculated. Only genes for which the 25^th^ percentile of expression in the tissue type of interest was higher than the 90^th^ percentile of expression in all other tissues were considered as tissue specific genes. These steps were repeated for each tissue type. Note that tissue specific genes might change in the future when new datasets will be added.

Expression profiles of samples in the Human Eye Transcriptome Atlas were validated using the top 100 most specific cell type marker genes for retinal microglia, rods and cones, as determined by single cell RNA sequencing ^22^. T-distributed stochastic neighbor embedding (t-SNE) plots ^23^ were performed to analyze clustering of all 100 samples in the Human Eye Transcriptome Atlas based on the above mentioned cell type marker.

## Data availability

The sequencing raw data are available in the Gene Expression Omnibus Database under the following accession numbers: GSE148387 (conjunctiva and conjunctival melanoma), GSE149004 (conjunctiva, conjunctival carcinoma and papilloma), GSE155776 (pterygium), GSE164192 (cornea), GSE164193 (eyelid), GSE159358 (lacrimal gland), GSE164194 (optic nerve), GSE159357 (retina), GSE146887 (choroid/RPE and CNV membranes) and GSE147657 (hyalocytes). Processed sequencing data can easily be explored without bioinformatics expertise using the Human Eye Transcriptome Atlas website available at https://www.eye-transcriptome.com, -.de, -.org, www.eye-rna-seq.com, -.de, -.org, www.eye-map.org, www.ophthalmo-transcriptome.com, -.de, -.org, www.ophthalmo-rna-seq.com, -.de, -.org and www.ophthalmo-map.com, -.de, -.org, which all lead to the main domain at https://www.eye-transcriptome.com. The datasets section of the website also provides download links for the sequencing raw data mentioned above as well as links to the associated publications from our group.

